# Lethal and sublethal effects of synthetic and bio-insecticides on *Trichogramma brassicae* parasitizing *Tuta absoluta*

**DOI:** 10.1101/2020.11.20.391003

**Authors:** Zahra Nozad-Bonab, Mir Jalil Hejazi, Shahzad Iranipour, Mehdi Arzanlou, Antonio Biondi

**Author notes:** Corresponding author (ZN).

## Abstract

The invasive tomato leaf miner (TLM), *Tuta absoluta* (Meyrick) is an invasive pest on tomatoes worldwide. The main control measure against the pest has been chemical insecticides, but the pest developed resistance to many chemical classes. So alternative methods, such as biological control agents, alone or combined to chemical compounds must be evaluated to validate their synergistic actions. In this study, both lethal (concentration-mortality response) and sublethal effects of three synthetic insecticides, the bioinsecticide spinosad, as well as the entomopathogenic fungus *Metarhizium anisopliae* (Metschnikoff) Sorokin were studied on *Trichogramma brassicae* Bezdenko within *T. absoluta* eggs. To assess the sublethal effects, the lethal concentration 25% (LC_25_) of chlorantraniliprole, spinosad, abamectin and indoxacarb and LC_50_ value of *M. anisopliae* was sprayed on eggs and then offered at three time intervals to the parasitoids. Fertility and other life table parameters of the individuals emerged from treated eggs were estimated. The results showed that indoxacarb showed the highest deleterious sublethal effects on *T. brassicae*. On the other hand, *M. anisopliae* was the safest treatment to combine to *Trichogramma* with no significant effect on some parameters. The lowest LC_50_ value for *T*. *brassicae* was obtained for chlorantraniliprole followed by spinosad. Synergistic effect was observed when *M. anisopliae* and *T. brassicae* used together. Hence, this will be a promising integration against *T. absoluta*.

## Introduction

The tomato leaf miner (TLM), *Tuta absoluta* (Meyrick) (Lep., Gelechiidae) is one of the most important pests of tomatoes worldwide [1]. Origin of this pest is from South America, but it spread rapidly through continents within 5-10 years. The first report of this pest out of Latin America was in Spain in 2006 [2]. Then *T. absoluta* reported from Africa and Asia. It invaded African tomato fields in 2016 and spread in almost 54 countries causing 50-100 % crop loss in tomato and some other products. Distribution of *T. absoluta* was documented that is related to temperature and moisture. It is also adapted to cold, warm, wet or dry environments, of different African countries [3]. Based on Han, et al. [4] review, this pest because of it’s ability to adapt to newly invaded area, high reproduction potential and … could destroy the tomato fields and greenhouse in Asian countries like Iran, after first report in Turkey in 2009. This pest invaded Iran’s fields in 2009 possibly via Turkey or Iraq borders [5].

This is a serious pest on some solanaceous crops having tomato as main host plant [6, 7]. According to Biondi et al. [1], potato and European black nightshade (*Solanum nigrum*) are suitable hosts for this pest. Furthermore, *T. absoluta* can oviposit and feed on some species of Amaranthaceae, Convolvulaceae, Fabaceae, and Malvaceae. Female needs to contact with oviposition stimulates of the host plants to begin ovipositing. The female can lay up to 260 eggs. The larvae are leaf miners and feed on leaf mesophyll between two epidermis. This behavior, protects larvae against non-systemic insecticides; moreover, this pest is well known to have developed resistance to various insecticide classes [8]. Because of this feeding behavior, short development time, several number of generations and rapid adaptation to different ecological conditions, this pest has been a most problematic and serious insect pest of tomatoes and can cause 100% damage if no management measure was adopted [1]. On the other hand, the use of insecticides in crop systems causes undesirable effects such as developing resistance and deleterious effects on beneficial arthropods [9–11]. In this context, obvious choices in sustainable pest management systems are biological control agents such as parasitoids, predators and entomopathogens [12–18]. Among the various parasitoid species parasitizing *T. absoluta* in both the native and the new invaded range [1, 19, 20]. Idiobiont oligophagous species of *Trichogramma* have shown to have a good potential to be employed in integrated pest management (IPM) programs [21–24]. Few investigations are dealing with integration of *Trichogramma* spp. with other control measures such as predators, entomopathogens and pheromone traps to control *T. absoluta* in Iran. These studies showed high importance of these egg parasitoids in IPM programs of *T. absoluta* [25–27]. Wide range of tolerance to environmental changes, easy method of rearing, killing their hosts prior to damage and a measurable host preference despite of a wide host range are advantages that make *Trichogramma* spp. valuable biological control agents [21, 28]. The host species may dramatically affect the fitness of *Trichogramma* parasitoids, such as size, longevity, fecundity and host acceptance [23, 29–31]. Some Trichogrammatidae species, such as *T. achaeae* (Nagaraja & Nagarkatti) [22, 32], *T. euproctidis* (Girault) [33, 24], and *T. evanescens* Westwood [34], have been reported as effective egg parasitoids of *T*. *absoluta*. Unfortunately, none of these species is still extensively used commercially. In Iran, *T. brassicae* Bezdenko (Hym.: Trichogrammatidae) is the only species that is reared in restricted scale and thus have potential for future implication in IPM programs in Iran.

Other important agents in biological control of *T. absoluta* are pathogens which are used commercially nowadays. Among pathogens, fungi are promising due to diversity and adaptation to agroecosystems [35]. Most of the commercially used fungal entomopathogens are belonging to the genera of *Metarhizium*, *Beauveria*, *Lecanicillium* and *Isaria*. They can directly penetrate into the insect cuticle and are able to cause epizootics. Also they can be easily produced in large quantities [36, 37]. *Metarhizium anisopliae* (Metschnikoff) Sorokin (Hypocreales: Clavicipitaceae) is a well-known entomopathogen belongs to Pezizomycotina: Sordariomycetes [38]. Some members of this fungi are known as saprophytes and some species or sub-species are registered as entomopathogen [35]. Sometimes *Metarhizium* spp. produce sclerotia in cultural media which makes this fungus resistant to harsh conditions of cold seasons [39]. There are some reports of high virulence of this fungus on *T. absoluta* eggs, larvae, pupae and adults [13, 40, 41]. *Metarhizium anisopliae* is a promising entomopathogenic fungus which can be used in commercial scale [39]. Therefore, a suitable integration with other control tools is needed to provide effective and sustainable control.

According to Suh, et al. [42], spinosad and prophenofos had high toxic effects on *T. exiguum* Pinto & Platner in *Helicoverpa zea* (Boddie) eggs and reduced its longevity and hatch rate. Their results documented that spinosad was effective even after four days and had significant side effects on parasitoids. Spinosad, as a bioinsecticide is used both in traditional and organic cultures, therefore studying on side effects of spinosad weather on pest or biological control agents is so important. In a literature review, Biondi, et al. [43] found that spinosad is more toxic for Hymenoptera than other parasitoid orders, although it is more selective to bees. Moreover, modern insecticides can be seem to be safe, while residue of them can affect fecundity, longevity and sex ratio of biological control agents. Hence, a comprehensive evaluation of insecticide effects should be done by considering sublethal effects on natural enemies as well [9, 44, 45]. Therefore, combining insecticides and parasitoids may not have the expected results.

Because compounds used against *T. absoluta*, affect directly and/or indirectly its natural enemies, evaluating such effects is necessary for choosing suitable insecticides in tomato leaf miner IPM programs. This study was conducted to evaluate lethal and sublethal effects of some insecticides and *M. anisopliae* on *T. brassicae*, the prevalent egg parasitoid species of *Trichogramma* in Iran. The selected insecticides were the most effective ones among compounds tested on *T. absoluta* based on a previous study [46].

## Materials and methods

### Rearing *Tuta absoluta*

Different larval instars of *T. absoluta* were collected from a damaged tomato field in Bilasuvar County (39 ° 39ʹ 31.37ʺN 48 ° 34ʹ 67.06ʺE) in Ardabil province in Northwest of Iran and moved to a greenhouse section of the Department of Plant Protection, University of Tabriz, Tabriz, Iran. The larvae were kept at 27 ± 2 °C, 50 ± 10 % RH and 16: 8 h (L: D) photoperiod and fed on foliage of greenhouse grown tomato plants and maintained until adult emergence. The adults then were transferred to 80 × 70 × 60 cm wooden cages covered with organdy cloth within which two or three potted tomato plants (20 - 30 cm height) were included and let them to mate and lay their eggs on the plants for 24 hours. The adults were fed with 10% sugar solution (renewed every three days). After 24 h the plants were shaken within the cage to remove the moths from them. Twenty four hour old eggs were used in bioassays.

### *Trichogramma brassicae* rearing

The egg parasitoid *T. brassicae* was provided from a private insectarium in Parsabad, Ardabil province. The stock culture of the parasitoid was reared on *Ephestia kuehniella* (Zeller) (Lep.: Pyralidae) eggs within glass tubes (1cm diameter, 6 cm length) in a growth chamber at 27 ±1°C, 50±5 % RH and 16:8 h (L:D) photoperiod. These parasitoids were reared on *T. absoluta* eggs for two further generations, prior to using in experiments.

### *Metarhizium anisopliae* cultures

Source of the entomopathogenic fungus *M. anisopliae*, was provided by the laboratory of Biological Control of Insects, University of Tehran. The fungus was cultured on potato dextrose agar (PDA) medium in Petri-dishes (9 cm in diameter) at 25±1 °C. Ten days later, the cultures with well-developed spores were washed with distilled water + 0.2 % surfactant Tween-80^®^. After filtering the spore suspension by glass wool, the number of spores was counted using a haemocytometer (Assistent^®^).

### Lethal effects of insecticides on *Trichogramma brassicae*

Based on a previous work (Nozad-Bonab, et al., 2017), four chemical insecticides (spinosad (Laser^®^ 48 SC), indoxacarb (Steward^®^ 30 WG), abamectin (Vertimec^®^ 1.8 EC) and chlorantraniliprole (Coragen^®^ 18.5 SC) were chosen as effective insecticides for *T. absoluta* control.

The lethal effects of above insecticides were studied on *T. brassicae*. To assess lethal effects, 120 tomato leaf miner eggs were put on a piece of paper (1.5 × 3 cm) in a glass tube (6 cm length, 1cm diameter) and exposed to *T. brassicae* females. Five days later when blackhead stage of parasitized eggs appeared, these were sprayed by those insecticides using a Potter spray tower (Burcard Scientific ^®^) (5 ml insecticide solution under 0.5 bar pressure). The ranges of concentrations were 0.09 – 1.26; 0.6 – 4.5; 0.55 – 11.1; 0.024 - 0.72 mg a.i. L-1 of abamectin, indoxacarb, chlorantraniliprole, spinosad, respectively. Tween-80^®^ was used as surfactant at a concentration of 0.05 % (v/v) in all treatments. In the control the eggs were sprayed with distilled water + Tween-80^®^. Five days later the number of emerged parasitoid was recorded and the LC_50_ values were estimated. The experiment had three replicates with 40 insects each. The mortalities were corrected using Abbott’s formula [47] and the LC_50_ values were estimated using the probit procedure of SPSS [48].

### Sublethal effects of insecticides on *Trichogramma brassicae*

For evaluation of sublethal effects of the above mentioned insecticides as well as entomopathogenic fungus, *M. anisopliae* on *T. brassicae*, the LC_25_ values of chemical insecticides and the LC_50_ value of entomopathogenic fungus (0.07, 1.82, 1.44 and 0.056 mg ai/l of spinosad, indoxacarb, abamectin and chlorantraniliprole respectively, and 10^4^ spore/ml of the fungus; obtained in a previous work, [46], were sprayed on 50 *T. absoluta* eggs upon a piece of paper (3 cm length, 1.5 cm width). The treated eggs were exposed to the parasitoids 0, 24 and 48 h later. Thirty couples of the parasitoids were selected and transferred in glass tubes (6 cm length, 1cm diameter). Very small honey droplets (20%) were placed on a piece of paper (2 cm length, 1 cm width) and deposited in tubes. All the tubes were kept in a growth chamber (27 ± 1 ºC, 60 ± 10 % RH and 16: 8 h photoperiod) until the end of the study. Development time, emergence rate, longevity and fecundity of the progeny, were thus assessed.

The range of spore concentration of the *M. anisopliae* was determined as 1.2 × 10^2^ – 1.2 × 10^6^ spore/l. Fifty *T. absoluta* eggs were sprayed by LC_50_ of *M. anisopliae* by using above mentioned Potter spray tower and exposed to *T. brassicae* after drying in room condition. The experiment was repeated three times at different days.

### Combination of *Trichogramma brassicae* and insecticides or entomopathogenic *Metarhizium anisopliae*

In other experiment, 20 eggs of *T. absoluta* treated by LC_25_ of insecticides or LC_50_ of *M. anisopliae* by using a Patter Spray Tower, exposed to *T. brassicae* females immediately after drying, 24h or 48 h after incubation in laboratory condition, and then they were kept in a growth chamber until larvae emergence. This experiment was repeated for 30 parasitoids. The number of *T. brassicae* adults and *T. absoluta* larvae were counted at the end of the experiment and mortality rate was estimated as number of parasitized eggs to available ones. This study had three sets of control treatments, 1. The untreated eggs on leaflet to ensure their healthy, 2. The parasitized eggs without insecticides to ensure the successful parasitism and 3. The treated eggs by insecticides and *M. anisopliae* for ensuring the effectiveness of insecticides and entomopathogen. Each experiment had four replications.

### Data Analysis

Life table parameters, including gross reproductive rate (GRR), net reproductive rate (R_0_), intrinsic rate of increase (r_m_), finite rate of increase (λ), generation time (T), doubling time (DT), intrinsic birth rate (b) and intrinsic death rate (d), were estimated according to Carey [49] and Biondi, et al. [44]. Variance of the parameters was estimated using Jackknife pseudovalues. One-Way ANOVA and post hoc test of Tukey (α=0.05) were used to compare means of the treatments. To categorize the binary relationship between different control agents, in antagonistic, additive or synergistic category, a modified method of Koppenhöfer and Kaya [50] by Yii, et al. [51] was adopted. This method is based on testing discrepancy of the observed mortality from an expected mortality calculated as ME = MC + MB (1 – MC/100) by using a chi-square test (df = 1), where ME, MC and MB are expected mortality, mortality by chemical factor and mortality by biological factor, respectively.

In chi-square test, χ^2^ = (MCB – ME)2/ME, where MCB is the observed mortality for the parasitoid–insecticide combinations. If the calculated χ^2^ value exceeds the critical value of the Chi square table (3.84, df=1), implies non-additive (synergistic or antagonistic) relation between the two agents [52]; otherwise it is an additive relation. In circumstance which, null hypothesis of additive relation was rejected, if the D = MCB – ME is a positive value, then the relation is considered a synergistic type, nonetheless it is an antagonistic one.

## Results

### Lethal effects

The LC_50_ values estimated for the examined insecticides on *T. brassicae* are shown in table 1. The results indicated that chlorantraniliprole had the lowest LC_50_ value, followed by spinosad, indoxacarb and abamectin. The LC_50_ value on *T. brassicae* was 5.33 times higher than that of the *T. absoluta* in our previous study [46]. Moreover, Indoxacarb was 2.975 times more toxic on the parasitoid than tomato leaf miner. While spinosad and abamectin had almost similar toxic effect both on host and parasitoid. The LC_50_ values of spinosad, abamectin, indoxacarb and chlorantraniliprole on *T. absoluta* (0.14, 3.61, 3.99 and 0.11 mg ai./l, respectively) were estimated by Nozad-Bonab, et al. [46].

**Table 1.**
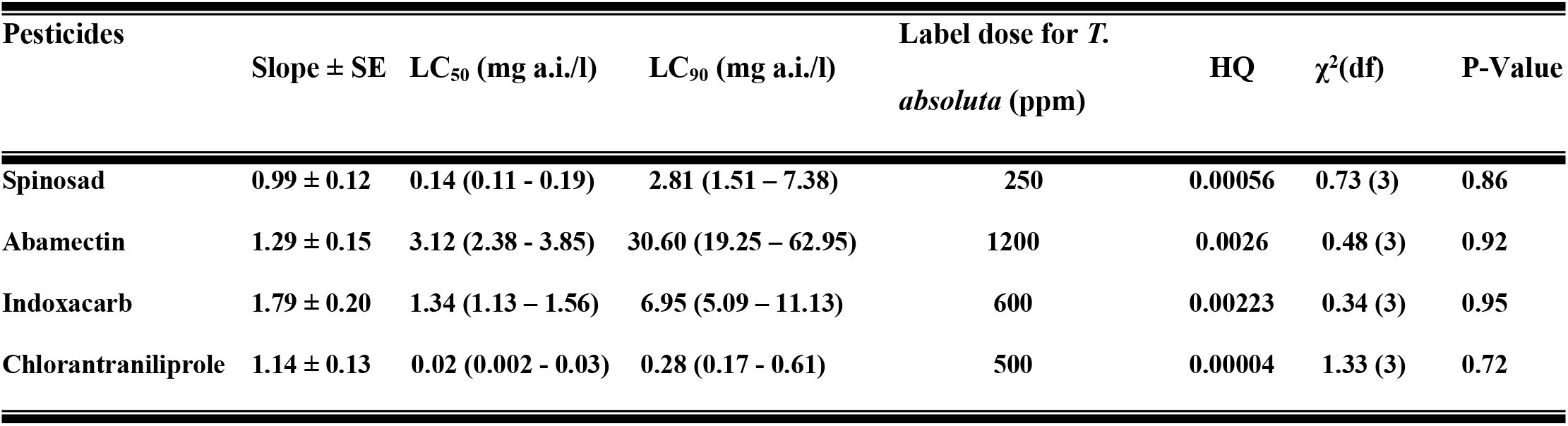
Summary of probit analysis results and estimated Lethal Concentrations (LC_50_ and LC_90_) of the chemical insecticides tested on *Trichogramma brassicae* juvenile stages within *Tuta absoluta* eggs.

| Pesticides | Slope ± SE | LC<sub>50</sub> (mg a.i./l) | LC<sub>90</sub> (mg a.i./l) | Label dose for *T. absoluta* (ppm) | HQ | χ<sup>2</sup>(df) | P-Value |
| --- | --- | --- | --- | --- | --- | --- | --- |
| Spinosad | 0.99 ± 0.12 | 0.14 (0.11 - 0.19) | 2.81 (1.51 – 7.38) | 250 | 0.00056 | 0.73 (3) | 0.86 |
| Abamectin | 1.29 ± 0.15 | 3.12 (2.38 - 3.85) | 30.60 (19.25 – 62.95) | 1200 | 0.0026 | 0.48 (3) | 0.92 |
| Indoxacarb | 1.79 ± 0.20 | 1.34 (1.13 – 1.56) | 6.95 (5.09 – 11.13) | 600 | 0.00223 | 0.34 (3) | 0.95 |
| Chlorantraniliprole | 1.14 ± 0.13 | 0.02 (0.002 - 0.03) | 0.28 (0.17 - 0.61) | 500 | 0.00004 | 1.33 (3) | 0.72 |

### Sublethal effects

Longevity was 3.32 days in control and 1.32, 1.47, 1.54, 1.32 and 3.06 d, in adult parasitoids that emerged from *T. absoluta* eggs sprayed with abamectin, chlorantraniliprole, spinosad, indoxacarb and *M. anisopliae*, respectively. The chemical insecticides caused significant effects on life table parameters in comparison with control (Table 2). On the other hand *M. anisopliae* had a moderate and delayed effect on biostatistics of *T. brassicae*. For example effect on *r*_*m*_ *b*, λ was observed only after 24h. Also some parameters such as GRR, DT and T did not affect by *Metarhizium anisopliae*.

**Table 2.**
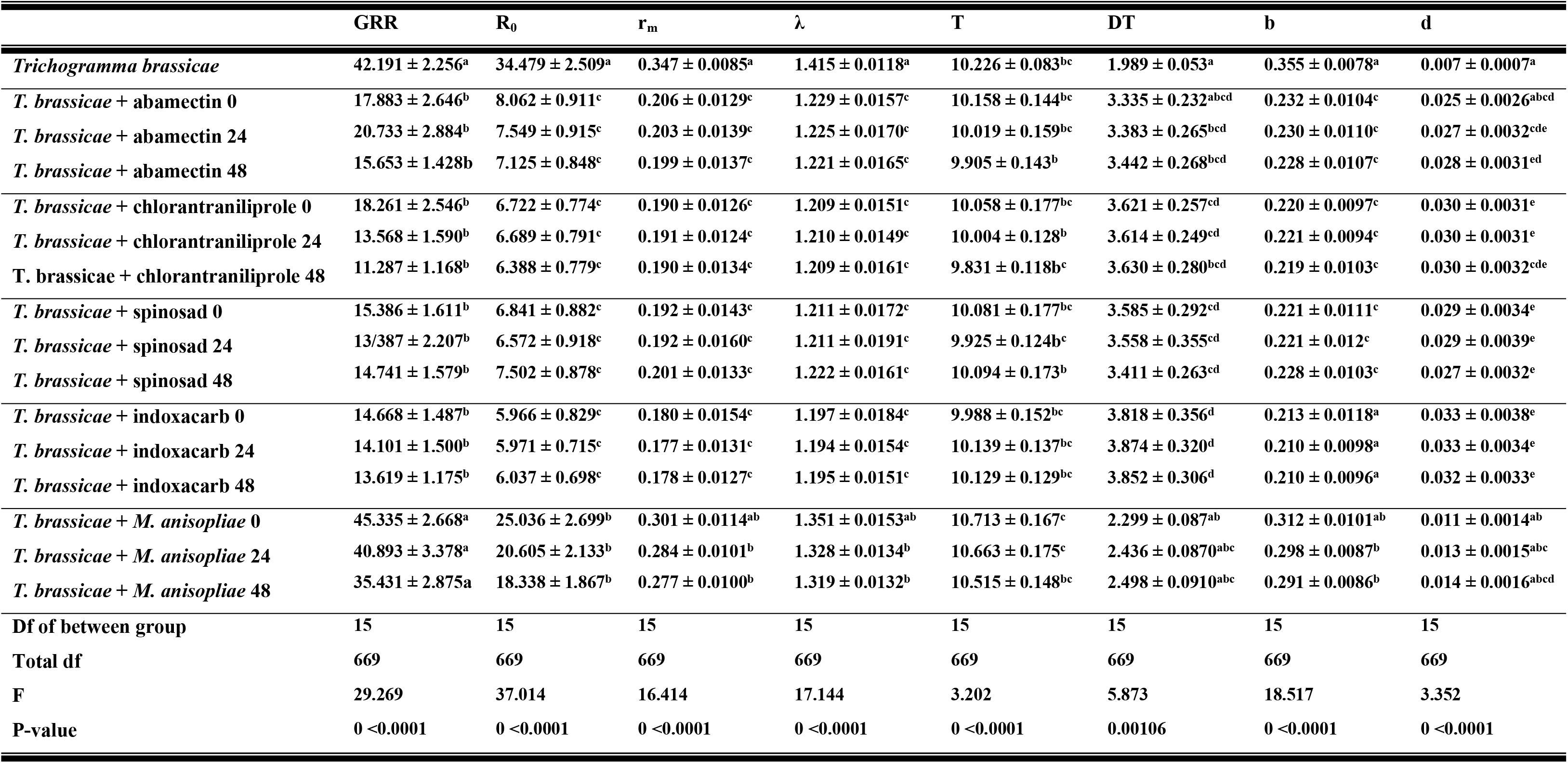
Life table parameters (means ± SE) estimated for *Trichogramma brassicae* developed on *Tuta absoluta* eggs sprayed with Lethal Concentration 25% (LC_25_) of abamectin, chlorantraniliprole, spinosad, indoxacarb and *Metarhizium anisopliae* 0, 24 and 48 hours prior parasitism.

Chemical insecticides changed the gross fecundity rate compared to control. However, no significant interaction was observed between insecticides and exposure times. In addition, the entomopathogenic fungus *M. anisopliae* had no significant time-dependent effect. Nevertheless, net reproduction rate as well as birth rate showed significant difference with control in the parasitoid and pathogen integration. It seems that in contrast to chemical insecticides, entomopathogenic fungus had no effect on intrinsic rate of increase. Sublethal effects of the tested insecticides was not significant on generation time, but doubling time was longer for indoxacarb. Birth rate and death rate was respectively maximum and minimum in control. The entomopathogenic fungus *M. anisopliae* developed and killed the eggs, and the survived eggs were so low quality that adversely affected the parasitoid preference, but this negative effect was less than insecticides.

Although insecticides and entomopathogen had some negative effects on parasitoid, but the combination of them could increase the *T. absoluta* eggs mortality. According to results of table 3, the combination of *M. anisopliae* and *T. brassicae* had a synergistic effect in simultaneous applications. Other combinations showed rather additive effects except abamectin + parasitoid and chlorantraniliprole + parasitoid that were mainly antagonistic. Perhaps, longer developmental time of the infected host extends available time for parasitism, thus combination of these agents can be more effective than the insecticides + the parasitoid.

**Table 3.** The mortality rate of *Tuta absoluta* in an integrated system including the parasitoid *Trichogramma brassicae* + each one of the entomopathogenic fungus, *Metarhizium anisopliae*, or chemical insecticides, spinosad, abamectin, indoxacarb and chlorantraniliprole

| Combination | Time (h) of the parasitoid | % observed mortality | % expected mortality | $\chi^2(df)$ | Type of effect |
| --- | --- | --- | --- | --- | --- |
| Spinosad + parasitoid | 0 | 74.47 | 71.11 | 0.16 (1) | additive |
|  | 24 | 77.54 | 68.84 | 1.10 (1) | additive |
|  | 48 | 78.39 | 68.18 | 1.53 (1) | additive |
| Abamectin + parasitoid | 0 | 68.82 | 74.15 | 0.38 (1) | antagonistic |
|  | 24 | 70.52 | 70.48 | 0.28 (1) | additive |
|  | 48 | 74.53 | 67.32 | 0.77 (1) | additive |
| Indoxacarb + parasitoid | 0 | 70.46 | 69.59 | 0.01 (1) | additive |
|  | 24 | 72.03 | 68.02 | 0.24 (1) | additive |
|  | 48 | 71.68 | 69.90 | 0.04 (1) | additive |
| Chlorantraniliprole + parasitoid | 0 | 73.25 | 73.39 | 0.0003 (1) | antagonistic |
|  | 24 | 74.10 | 68.84 | 0.40 (1) | additive |
|  | 48 | 74.84 | 67.32 | 0.84 (1) | additive |
| *Metarhizium anisopliae* + parasitoid | 0 | 94.30 | 76.44 | 4.17 (1) | synergistic |
|  | 24 | 94.30 | 84 | 1.26 (1) | additive |
|  | 48 | 95 | 84 | 1.44 (1) | additive |

## Discussion

Biological control agents are often more sensitive to insecticides than targeted pests. It is may be due to shorter exposure time to insecticides and lower doses that they receive [53]. Since mechanism of physiological selectivity of spinosad is unknown, researchers cannot explain why resistance to spinosad has evolved by insects. However low penetration rate into integument, change in site of action or increased rate of the insecticide metabolism are possible explanations for spinosad selectivity for wasps. Fernandes et al. [54] suggested spinosad as a moderately toxic compound on Vespidae and Apidae. They explained that low penetration rate is because of cohesion with the integument and large molecular weight of spinosad. In our study, no selectivity was observed between the parasitoid *T. brassicae* and its host *T. absoluta*. Also Suh et al. [42] evaluated spinosad as a toxic compound to *Trichogramma exiguum*. But Hewa-Kapuge et al. [55] reported indoxacarb as a low toxicant compound on *Trichogramma* nr. *brassicae* in laboratory conditions while it reduced adults emergence in field. Also they reported emamectin as a moderately toxic insecticide against parasitoids. The difference observed between our study and those of the Hewa-Kapuge et al. [55] is partly due to different egg shell characteristics of different hosts used in these studies. In their study *H. armigera* with a thicker egg shell was used as host.

In addition to the lethal effects, the sublethal effects and life table parameters can also determine the degree of toxicity of a pesticide to natural enemies and pests. So that, Lundgren and Heimpel [56] documented that the longevity of *T. brassicae* was 4 and 2 days with and without feeding on honey. Also Orr et al. [21] also recorded longevity to be 4 days for *T. brassicae.* Both studies partially agreed our study. On the other hand, Afshari et al. [57], showed that indoxacarb reduced longevity and efficiency of *T. brassicae*. Suh et al. [42] also reported that spinosad reduced emergence rate and longevity of *T. exiguum* on *Helicoverpa armigera* (Boddie) eggs. But spinosad did not change fecundity, sex ratio or frequency of brachyptery. Spinosad was classified as moderate toxic on *T. exiguum* females. It seems that spinosad could not penetrate in the host egg and the parasitoid adults affected by spinosad only at emergence. Difference in parasitoid species and the insecticides’ dose may partly explain differences in our results with this study.

There is another opinion about spinosad, Medina et al. [58] showed that first and second larval stages of *Hyposoter didymator* (Thunberg) (Hym., Ichneumonidae) were affected less than the third instar larvae within body of host, larva of *Spodoptera littoralis* (Boisduval) (Lep.: Noctuidae). Since the 1^st^ and the 2^nd^ instar larvae feed the host hemolymph, they gain the lower spinosad residue than the 3^rd^ instar larvae that feed the cellular tissue. Also adults are more highly affected in oral rather than contact exposure. The adults chew the silken cocoon and in this way they can intake some residue of spinosad and die. They reported that spinosad had higher oral proportional to contact toxicity for this parasitoid because the thick cuticle of the host act as a preventive barrier. Undesirable effects of spinosad on *T. brassicae* life table parameters, may be due to the mentioned reasons. It depends on both host and natural enemy species that which one of the contact or oral effects of spinosad is stronger. Ruiz et al. [59] indicated that the contact effect of spinosad is more than oral effect on *Diachasmimorpha longicaudata* (Ashmead) (Hym.: Braconidae). Furthermore, they found that sublethal doses of spinosad had deleterious effects on fecundity, survival and partly on sex ratio.

Fernandes et al. [60] documented that the reason of reduction in fecundity and size of parasitoids progeny may be due to accumulation of spinosad in ovaries. This can explain the reason of reduction of fecundity in our study. Similarly, Schneider et al. [61] reported lethal and sublethal effects of spinosad on *Hyposoter didymator* (Thunberg) (Hym.: Ichneumonidae). However, the mechanism of spinosad toxicity on parasitoid wasps is almost unknown. The parasitoids intake so few host egg chorion, that one may not expect detectable effects. Perhaps those amounts that penetrate into eggs are responsible for spinosad effects. Cônsoli et al. [62] reported that the effect of this insecticide on *Trichogramma galloi* Zucchi refer to its effect on neural system and especially nicotinic receptors that causes parasitoid paralyze. Also the female parasitoids are directly exposed to insecticides during egg probe and host feeding. Spinosad had deleterious effect on female at first day. All these theories can explain the sublethal effects of spinosad. Also, Hossain and Poehling [63] showed that, a part of the insecticides penetrate to underside layers of host eggs and larval skin and the remainder of them absorbed by host tissue, and fed by the parasitoid and in this way, parasitoid is exposed to sublethal doses of abamectin and spinosad. On the other hand, Blibech et al. [64] documented spinosad as a safer compound for three species of *Trichogramma* compared to deltamethrin. This is in contrast to our study that categorizes spinosad as similar to the other insecticides. They found spinosad as low to moderate toxic compound against these parasitoids and argued about difference among species response to different insecticides.

On the other side, Ahmadipour et al. [65] studied six populations of *Trichogramma* in Iran (Amol, Baboulsar, Mashhad, Langroud, Shiroud and Some-e Sara) on *T. absoluta* eggs in laboratory conditions. They reported significant difference between populations. The highest parasitism rate was 54 % in Baboulsar population. Also, there was no significant difference for egg parasitism on both sides of tomato leaves when egg density was the same on both sides. The tested *Trichogramma* population in our study was from Bilesovar with 33.6 percent parasitism on *T. absoluta* eggs which was close to Shiroud population (36.4 %) of Ahmadipour et al. [65].

On the other side, Abamectin not only has a high lethal effect on *T. absoluta* [46], but also, causes sublethal effects on biological control agents like *T. brassicae*, hence it should be used cautiously [44]. Undesirable sublethal effects of both abamectin and spinosad such as reduced fecundity and longevity, was reported on *Bracon nigrican*s Szépligeti (Hymenoptera: Braconidae). Their results agree our ones in presence of detectable sublethal effects on the life table parameters of parasitoid; *T. brassicae* in this case. On the other hand Hewa-Kapuge et al. [55] evaluated emamectin as a very toxic compound for *Trichogramma* nr. *brassicae*, but indoxacarb was safe for this parasitoid under laboratory conditions. However, indoxacarb had a toxic effect on *T.* nr. *brassicae* in field experiments may be due to high temperature in field. Also Wang et al. [66] observed adverse effect of abamectin on *T. nubilale* Ertle & Davis in their experiment. Cônsoli et al. [67] categorized the abamectin as moderate toxic on *T. pretiosum* which reduced adults emergence and parasitism. However, in our study toxicity of abamectin was in the range of the other insecticides. This is possibly due to effect of insecticides on oogenesis and developmental stages of the parasitoid that can affect the results. Also according to Carvalho et al. [68], residue of abamectin or spinosad on host egg chorion may cause adults mortality or reduce longevity and fecundity in *T. pretiosum*. On the other hand, these insecticides were slightly more toxic to female than male, which lead to sex ratio variation.

Since high amounts of compounds can be transferred from hemolymph to ovaries, spinosad or other insecticides may be traced in eggs and cause deleterious effects on next generation [58]. Medina et al. [58] documented that 55% of spinosad was appeared in ovaries of *Hyposoter didymator* (Thunberg) (Hym.: Ichneumonidae). Like the present study, results of Sattar et al. [69] appreciated spinosad and emamectin benzoate as a very toxic compound on *T. chiloni*s. In their study, indoxacarb reduced fecundity and was slightly harmful except on egg. Eventually sublethal doses of insecticides may affect behavior and physiological state of a parasitoid without a parallel increase in mortality and so create a new population equilibrium. Delpuech et al. [70, 71] reported weakening response of males to females by insecticide treatments. Consequently fecundity reduced and sex ratio became male-biased.

Despite the adverse effects of insecticides on parasitoids, the combination of them was additive and synergic. Ashraf khan [72] also documented that, the residual of insecticides like abamectin can reduce the emergence and parasitism of *T. chilonis*, but they can be used in integrated pest management with parasitoid. But Blibech et al. [64] were disagree with these results, they revealed that, the usage of deltamethrin and spinosad with *Trichogramma oleae, T. cacoeciae* and *T*. *bourarachae* was usefulness in olive tree ecosystem integrated pest management.

The main purpose of IPM studies is finding of the strategies for more biological control using and produce healthy food. Better knowledge of chemical insecticides, biological mortality factors and combining synergistic control measures can help for gain the promising results. Based on the results of this study, it seems that spinosad and abamectin were more toxic for *T. brassicae* in comparison with indoxacarb and chlorantraniliprole. Also the entomopatogen fungus was safe for parasitoid in sublethal dose, then *M. anisopliae* was more compatible compound with parasitoid compared with chemical insecticides. Entomopathogen fungus not only was safer for parasitoid but also, they showed synergistic effect in leaf miner eggs mortality in combination together. Nevertheless, these results need further experiments and require validation by field studies to gain better insights on *Trichogramma* species effectiveness for integrated control programs.

## Acknowledgments

The authors thank Dr. Masoud Taghizadeh for kindly providing the initial colony of *T. brassicae.* We also appreciate Miss Solmaz Khani for her great assistance in conducting experiments.

